# eUTOPIA: Solution for omics data preprocessing and analysis

**DOI:** 10.1101/462879

**Authors:** Veer Singh Marwah, Giovanni Scala, Pia Anneli Sofia Kinaret, Angela Serra, Harri Alenius, Vittorio Fortino, Dario Greco

## Abstract

Application of microarrays in omics technologies enables quantification of many biomolecules simultaneously. It is widely applied to observe the positive or negative effect on biomolecule activity in perturbed versus the steady state by quantitative comparison. Community resources, such as Bioconductor and CRAN, host tools based on R that have become standard for high-throughput analytics. However, there is a need for intuitive and easy-to-use platform to process omics data, visualize, and interpret results, which is computational skill neutral. We propose an integrated software solution, eUTOPIA, that implements a set of essential processing steps as a guided workflow presented to the user as an R Shiny application. eUTOPIA allows researchers to perform preprocessing and analysis of microarray data *via* a simple and intuitive graphical interface while using state of the art methods. eUTOPIA is free for academic use and can be obtained from the GitHub repository https://github.com/Greco-Lab/eUTOPIA.

## Introduction

Omics data have become an integral part of biological studies as researchers leverage these techniques to obtain a broader perspective of complex biological phenomena. Scientific community resources such as Bioconductor and CRAN [1] contain an extensive collection of computational tools developed in R programming [2] language to process omics data. Application of these tools, however, requires a deep understanding of the computation and statistical aspects of the methods employed, while further integration of these tools in a workflow requires a certain degree of proficiency in computer programming languages. We tasked ourselves with creating a solution by implementing “state of the art” practices to preprocess, statistically analyze, and visualize microarray data from multiple platforms. eUTOPIA’s workflow caters to the researchers with varied levels of computation and statistical skills. This solution fills a void in a research environment, thus allowing experimental biologists to process microarray data while avoiding critical errors due to lack of proficiency in programming languages and pipeline development. The guided interface simplifies the complex tasks to set up the study and allows for more focus on the biological interpretation of the results rather than the technical challenges of executing statistical methods and generating visual representations.

## Materials and methods

eUTOPIA is developed in R programming language with a graphical interface layer designed by using R Shiny [3] web development framework. The dynamic user interface allows the user to adjust the parameters and observe the effect *via* meaningful graphical representations at each step, thus enabling to make informed decisions. Currently, eUTOPIA is capable of processing data from four microarray platforms: Agilent gene expression two-color microarray data (Samples specific to different colors channels), Agilent gene expression one-color microarray data, Affymetrix gene expression microarray data, and Illumina methylation microarray data. A guided stepwise workflow is implemented to preprocess and perform a preliminary analysis. eUTOPIA’s workflow is provided in Fig 1.

**Fig 1.**
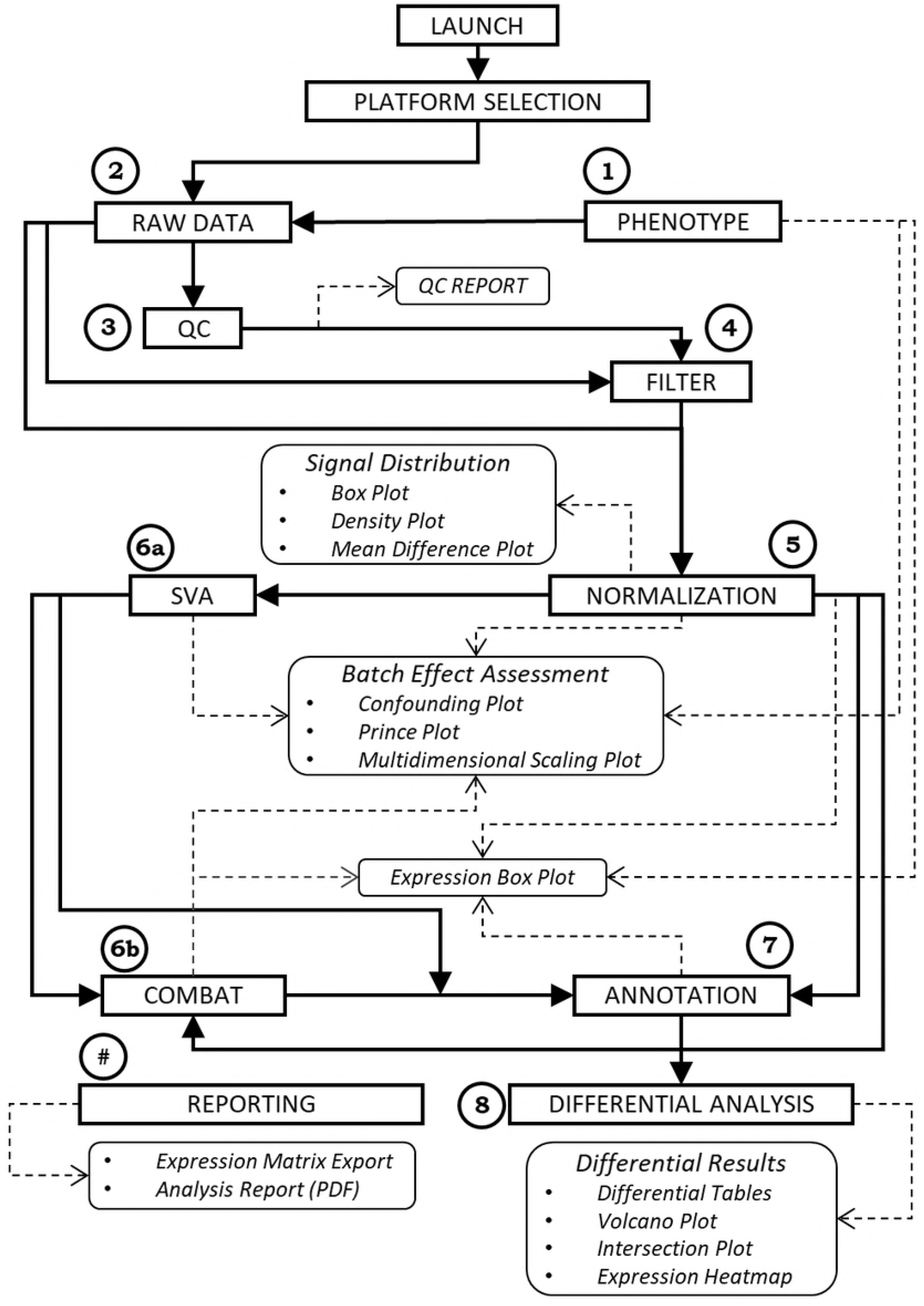
eUTOPIA workflow. Schematic representation of eUTOPIA’s guided workflow. The steps are represented by the rectangular box with sharp edges, and the step output is represented by the rectangular box with rounded edges. Analytical steps are numerically coded by circular labels from 1 through 8; an additional step labeled as ‘#’ is a reporting step with no specific order in the workflow. The labeled steps correspond to the steps of the analysis workflow in the eUTOPIA graphical user interface. The flexibility of defining alternate approaches for batch correction is captured in the workflow with the labeled steps 6a, and 6b that represent the subsections of BATCH CORRECTION step in the eUTOPIA analysis pipeline (S1 File). This workflow is adjusted according to the chosen data platform.

There are seven main steps in the workflow: ‘DATA INPUT’ (S1 File, p. 8-13), ‘QUALITY CONTROL’ & ‘FILTERING’ (S1 File, p. 14), ‘NORMALIZATION’ (S1 File, p. 15), ‘BATCH CORRECTION & ANNOTATION’ (S1 File, p. 20-29), and ‘DIFFERENTIAL ANALYSIS’ (S1 File, p. 30-31).

eUTOPIA requires that user provides a detailed phenotype information file (S2 File) with all biological and technical variables of the samples in the experiment.

A quality control report can be generated from microarray raw data by affyQCReport R package [4], yaqcaffy R package [5], arrayQualityMetrics R package [6], or shinyMethyl R package [7], depending on the microarray platform. It is essential that poor quality probes from the experimental data are omitted prior data normalization. In the gene expression specific platforms, this is accomplished by estimating the robustness of probe signals against the background (negative control probes). For Illumina methylation platforms, a detection p-value is computed by using the total DNA signal (methylated + unmethylated) against the background signal by minfi R package [8]. The expression value OR p-value threshold determined from the background signal is used to evaluate the probes across a percentage of sample specified by the user. Finally, the probes failing this evaluation are considered unreliable and thus filtered out.

Normalization of the expression and methylation signals distribution across the samples is performed respectively with methods from the limma [9] and minfi R packages. Methods *scale*, *quantile*, and *loess* perform normalization on log2-scaled intensities and ratios, and the *vsn* method uses a variance stabilizing transformation, which performs better for weakly expressed features. Normalization of Illumina methylation arrays is performed by using the methods from minfi R package with options of background subtraction, control feature measure, dye bias, and quantile normalization.

In microarray data analysis, a fundamental step is to attenuate the effects associated with technical variables (batch effects) while retaining the variation associated with biological variables. Batch effects can arise for multiple reasons, most commonly when the experiments are conducted in multiple batches, and the data is pooled together for processing. These batches can contribute to the variability of the features and could introduce a systematic error in their assessment, ultimately leading to incorrect results in the worst scenario [10]. Batch effects can be caused by known variables (*e.g.,* dye, RNA quality, experiment date, *etc*.) or by hidden effects not explained by the known variables. Known biological (*e.g.,* treatment, disease status, age, tissue, *etc*.) and technical (*e.g.,* dye, array, *etc*.) variables are provided by the user in the phenotype information, while unknown sources of variations can be identified by using the *sva* function from sva R package [11]. First, the impact of the technical variables is computed with *prince* function from swamp R package [12], and the correlation between both biological and technical variables is evaluated from the confounding plot which is generated by using the *confounding* function from the swamp R package. This information is used to identify batch variables as known technical or surrogate variables which are associated with strong sources of variation and are not correlated with biological variables of interest. These identified batch variables can be justifiably corrected to remove technical noise from the data. Finally, the correction is performed with *ComBat* [13] function from sva R package that employs an empirical Bayes approach to estimate systemic batch biases affecting many genes. The batch correction is carried out by specifying the variable of interest, any biological covariates, and a set of known batches or surrogate variables (obtained from the *sva* function described above). The batch correction process implemented in eUTOPIA applies the *ComBat* function iteratively to remove one batch covariate at a time while the rest are modeled as covariates of interest. The *ComBat* function can process only batch covariate at a time, and this process of blocking other batches ensures clear separation of variation adjustment effects for each batch without any interference from the adjustment of other batches.

Linear models allow to model the covariate dependencies between samples. The differential analysis is performed with linear model implementation in the R package limma. The *lmFit* function from the limma R package fits gene-wise linear models to the microarray data. The user defines the design for the model by providing the biological variable of interest and covariates (biological and technical batch variables). The contrasts of interest are then specified to obtain contrast specific coefficients from the original coefficients of the linear model. The *eBayes* function is applied to assess differential expression by using the fitted model with the contrast coefficients. Final reporting of the differentially expressed genes is performed by using the *toptable* function where adjusted p-value for the multiple comparisons can be obtained by specifying methods “Holm”, “Hochberg”, “Hommel”, “Bonferroni”, “Benjamini & Hochberg”, “Benjamini and Yekutieli” or “False Detection Rate”. Differential analysis results for comparisons defined by the user are reported in tabular format and with other meaningful visualizations. The dynamic plots help to perform a preliminary interpretation of the analysis results. The distribution of the differential features by fold-change magnitude and significance can be observed for a chosen contrast by means of the volcano plots. The expression profile of user-specified top significant features from one or more contrasts can be inspected from the heatmap. Comparison of differential features from different contrasts by set intersections is represented as Venn diagrams or UpSet plots. And the distribution of signal for one or more gene(s) of interest in sample annotations (*e.g.,* experimental condition) can be inspected by means of box plots. A user manual with sample data analysis and plot descriptions is provided in S1 File.

## Case study

The functional capabilities of eUTOPIA are showcased here by processing a publicly available dataset GSE92900 (Kinaret *et al.*, 2017) obtained from the GEO (Barrett *et al.*, 2013) repository. The phenotype table (S2 File) is used for sample annotations.

The quality control of the raw data is performed with the arrayQualityMetrics R package. The quality report represents the distribution of intensities in arrays, the comparison between arrays (Fig 2), individual array quality, and it also reports outliers. In Fig 2A, array 10 has high distance measures with the rest of the arrays as represented by the bright and dull yellow colored cells. This distance information can also be interpreted from the hierarchical cluster in the right margin of Fig 2A, where array 10 clusters separately from the others. Outliers are detected based on this distance information as arrays with exceptionally large distance to all other arrays. A summarized distance measure is determined for each array by summing up the distances to all other arrays, which is checked against an outlier threshold obtained on the basis of well-established interquartile range (IQR) rules defined by Tukey Fences. The outliers can be observed in Fig 2B, where array 10 has a large summarized distance from the rest of the arrays and can be markedly recognized as an outlier.

**Fig 2.**
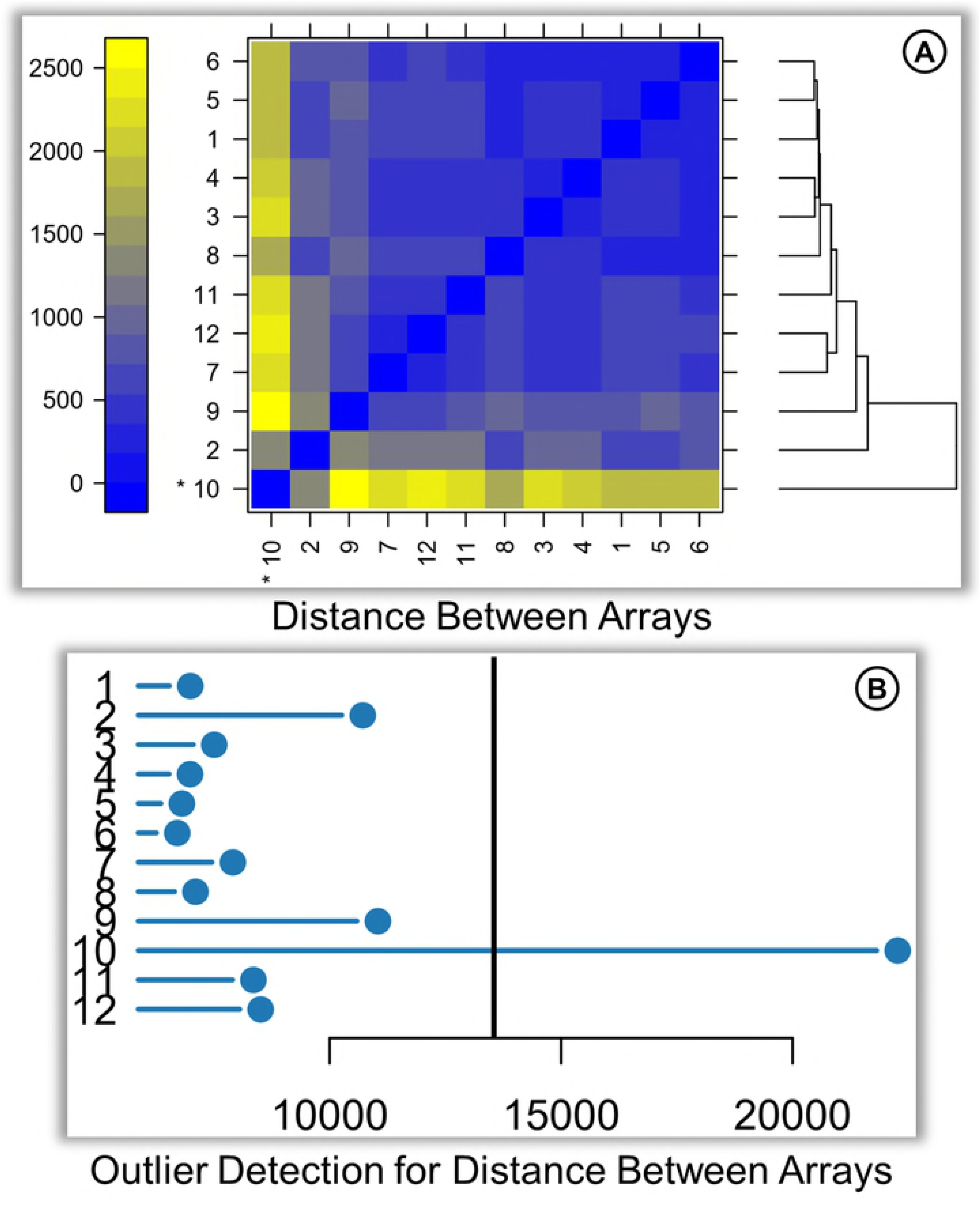
Sample QC. Subsection of arrayQualityMetrics quality control report. (A) Distance based on the mean absolute difference between array data represented as a Heatmap. The distance measure is represented by the color gradient from blue to yellow. (B) The pairwise distances for each array are summarized as a single representative value plotted on a horizontal bar chart. A threshold is determined based on the distribution of values across all arrays, represented by a vertical line. Arrays with the summarized distance larger than the threshold are considered outliers.

Normalization is performed by using the quantile method for between-array normalization. The normalized data can be observed by the representation of expression values as box plot, density plot, and mean difference plot (MDplot) (Fig 3).

**Fig 3.**
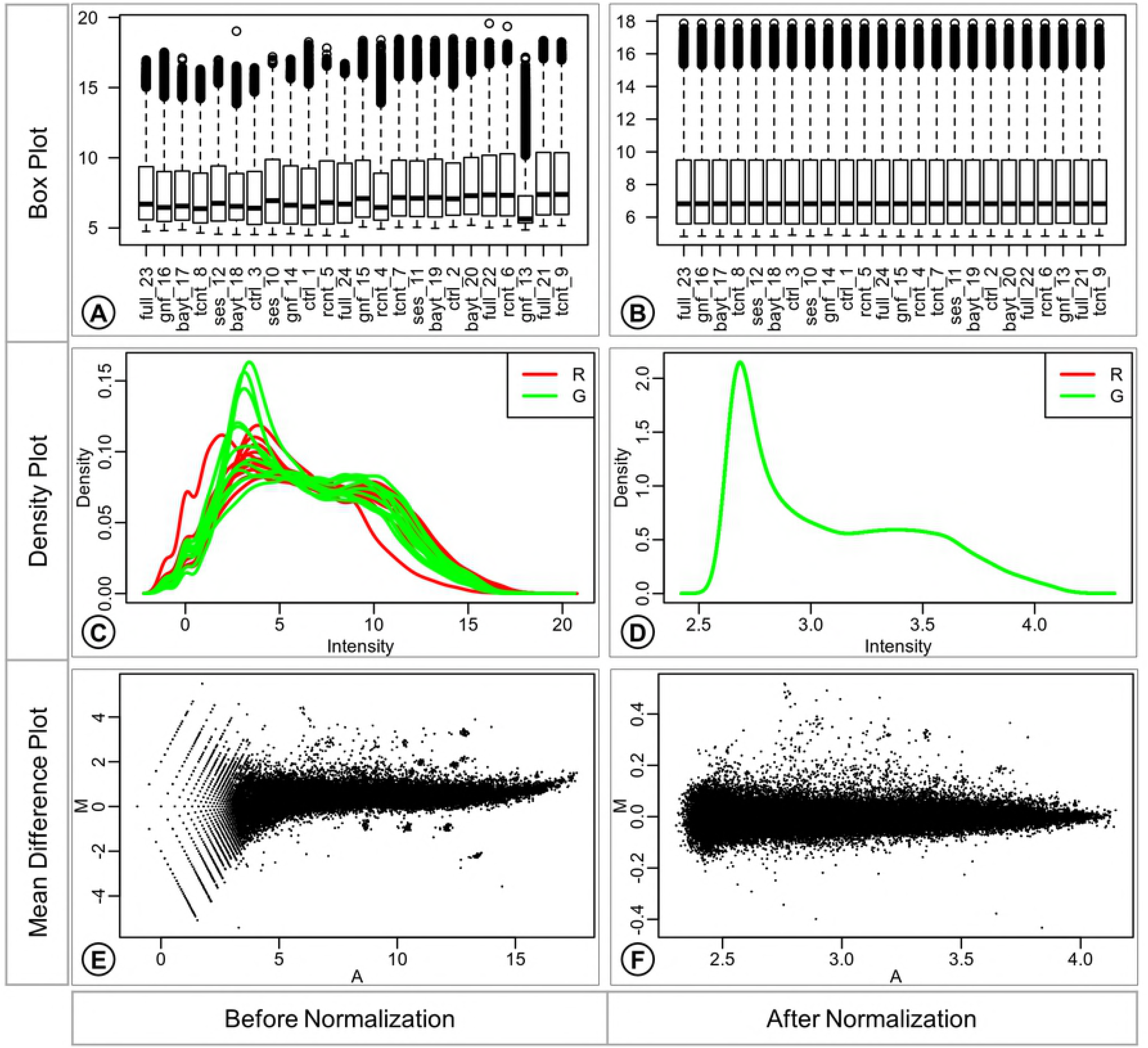
Normalization summary. Box plots (A; B), Density plots (C, D), and Mean-Difference plots (E, F) report expression values before and after normalization. The Box plots (A, B) report the distribution of expression values in each array. The Density plots (C, D) report the density distribution of the expression in individual color channels of each array. The expression intensity is plotted on the x-axis and the density values on the y-axis. The Mean-Difference plots (MD plot) (E, F) report agreement of the expression values between the individual channels of the array. The M-values (log-2 expression ratios) are plotted against the A-values (average log-2 expression values) as scatterplots. The A-values are plotted on the x-axis, and the M-values are plotted on the y-axis.

The difference in the distribution of expression values in different arrays can be observed in the box plot before normalization (Fig 3A), suggesting the need for adjustment of the distributions for fair comparison across the set of arrays. The distribution of expression values is harmonious across the arrays in the box plot after normalization with the quantile method (Fig 3B).

The difference in the distribution of expression values from individual channels of different arrays can be observed from the smoothed curves in the density plot (Fig 3C) before correction. Only a single smoothed curve is visible in the density plot after normalization (Fig 3D) since the channels from all arrays have the same distribution of expression values as a result of quantile normalization.

The larger scale of log-2 ratios (M-values on the y-axis) observed in the MDplot before normalization (Fig 3E) shows that the data points are farther away from zero log2 expression ratio, suggesting bias. The average log2 expression values also have a large scale (x-axis) before normalization. The smaller scale of M-value (y-axis) observed in the MDplot after normalization (Fig 3F) shows that data points are much closer to zero log2 expression ratios as the bias has been adjusted. From this plot, it is also possible to appreciate how the average log2 expression values have much smaller scale (x-axis) after normalization.

The known batches in the data are identified by inferring the variation associated with the sample annotations from the prince plot and by observing the significance of interrelatedness between the sample annotations from the confounding plot. The ‘group’ annotation variable is considered as the variable of interest. The prince plot (Fig 4A) displays that variables ‘slide’, ‘area’, ‘array’, ‘n.mice’, ‘operator’ and ‘date’ are associated with principal components representing high variation in the data. The confounding plot (Fig 4B) suggests that the variable ‘RIN’ is confounded with the variable of interest ‘group’ and thus cannot be taken forward for any data correction. Also, variables ‘slide’, ‘area’, ‘operator’, and ‘date’ are found to be confounded with ‘array’. The ‘array’ variable represents the highest variability in the data and is favored over ‘slide’, ‘area’, ‘operator’, and ‘date’ as a batch variable. Thus, it can be inferred from these plots that the final set of variables for batch correction is ‘array’, ‘dye’ (standard known batch), and ‘n.mice’ (Fig 4).

**Fig 4.**
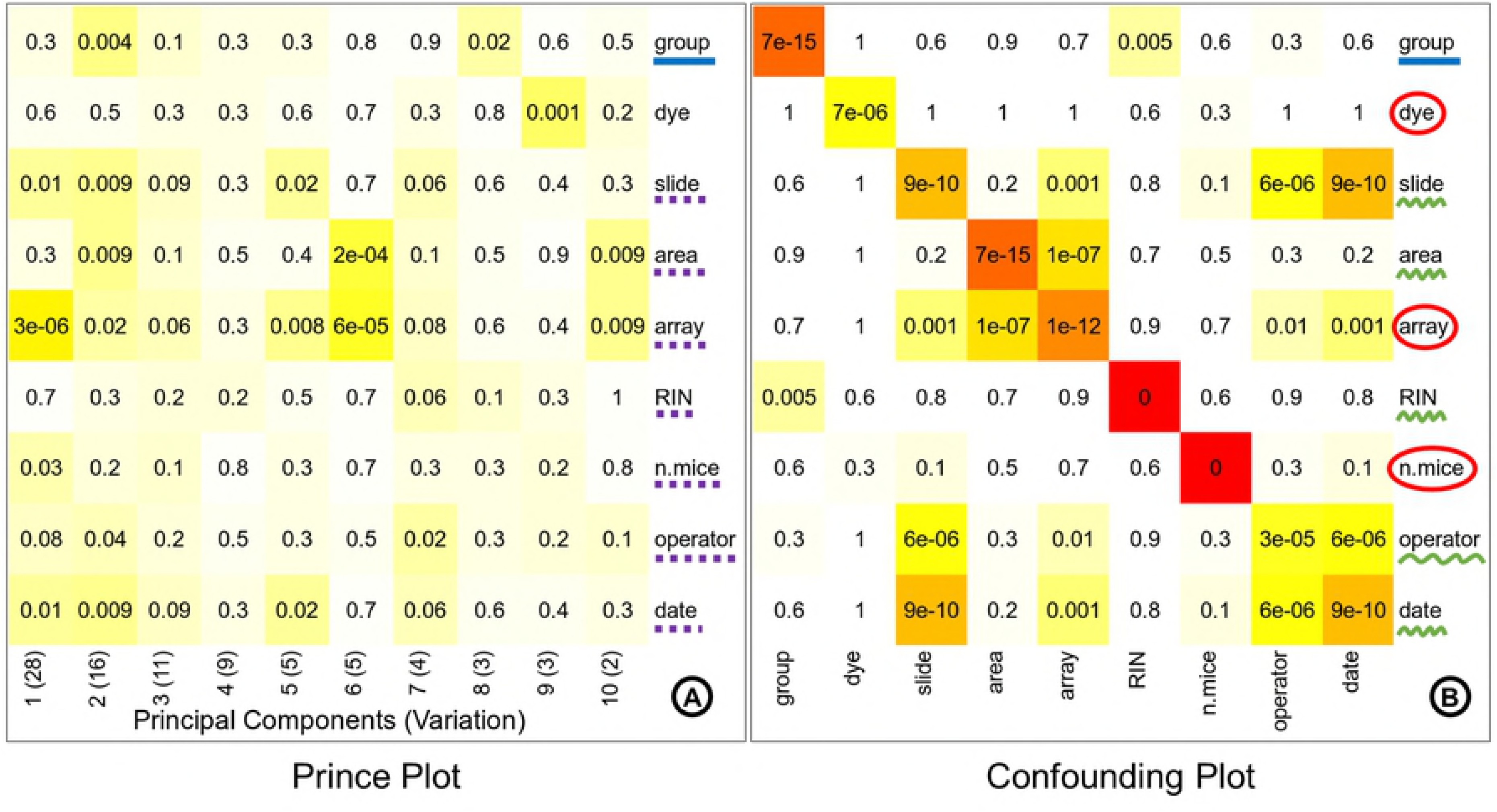
Identification of known batches. The prince plot (A) reports the association between the first ten principal components of the data and the annotation variables, principal components are represented in columns, and the sample annotations are represented in rows. The association p-value is represented in the cells by text labels, and background color in a chromatic scale. The row label underlined with solid blue line represents the variable of interest. The row labels underlined with dotted purple line represent other sources of high variation. The confounding plot (B) reports the association between all pairs of annotation variables; annotation variables are represented in both rows and columns. The cell text labels, cell background color, and row label notation with solid blue underline are same as the prince plot (A). The row labels underlined with squiggly green line represent the variables confounded with the variable of interest or other batch variables. The row labels circled by red outline are batch variables suitable for correction.

The effect of batch correction performed with eUTOPIA can be studied by comparing the variation information before and after correction. The first three principal components in the prince plot generated from data before batch correction (Fig 5A) represent 28%, 16%, and 11% variation in the data. The variable of interest ‘group’ was significantly associated (p-value: 0.004) with only the second principal component, while the identified batch variable ‘array’ was significantly associated with all three principal components (p-values: 3e-06, 0.02, and 0.06, respectively) and ‘n.mice’ was significantly associated with the first principal component (p-value: 0.03). In the prince plot generated from the data after batch correction (Fig 5B), the first three principal components represent 49%, 23%, and 11% variation in the data, which is a significant shift. The variable of interest ‘group’ is now observed to have a high association with all three principal components (p-values: 1e-13, 3e-14, and 7e-10, respectively). The batch variable ‘array’ has a comparably lesser significant association with the first principal component (p-value: 0.06) and batch variable ‘n.mice’ also has a comparably lesser association with the third principal component (p-value: 0.06). While batch variables ‘slide’, ‘area’, ‘operator’, ‘date’, and ‘dye’ are no longer associated with principal components representing high variation. Thus, allowing the user to check that the variation associated with the variable of interest is preserved while the noise associated with the batch variables has been corrected. The principal component analysis (PCA) plot generated from the data before batch correction (Fig 5C) displays samples scattered across the projected components with no obvious grouping of samples by the variable of interest. The PCA plot (Fig 5D) after the known batch correction displays a more discrete grouping of samples (smaller intragroup distances) and better separation of groups (intergroup distances) in the projected components.

**Fig 5.**
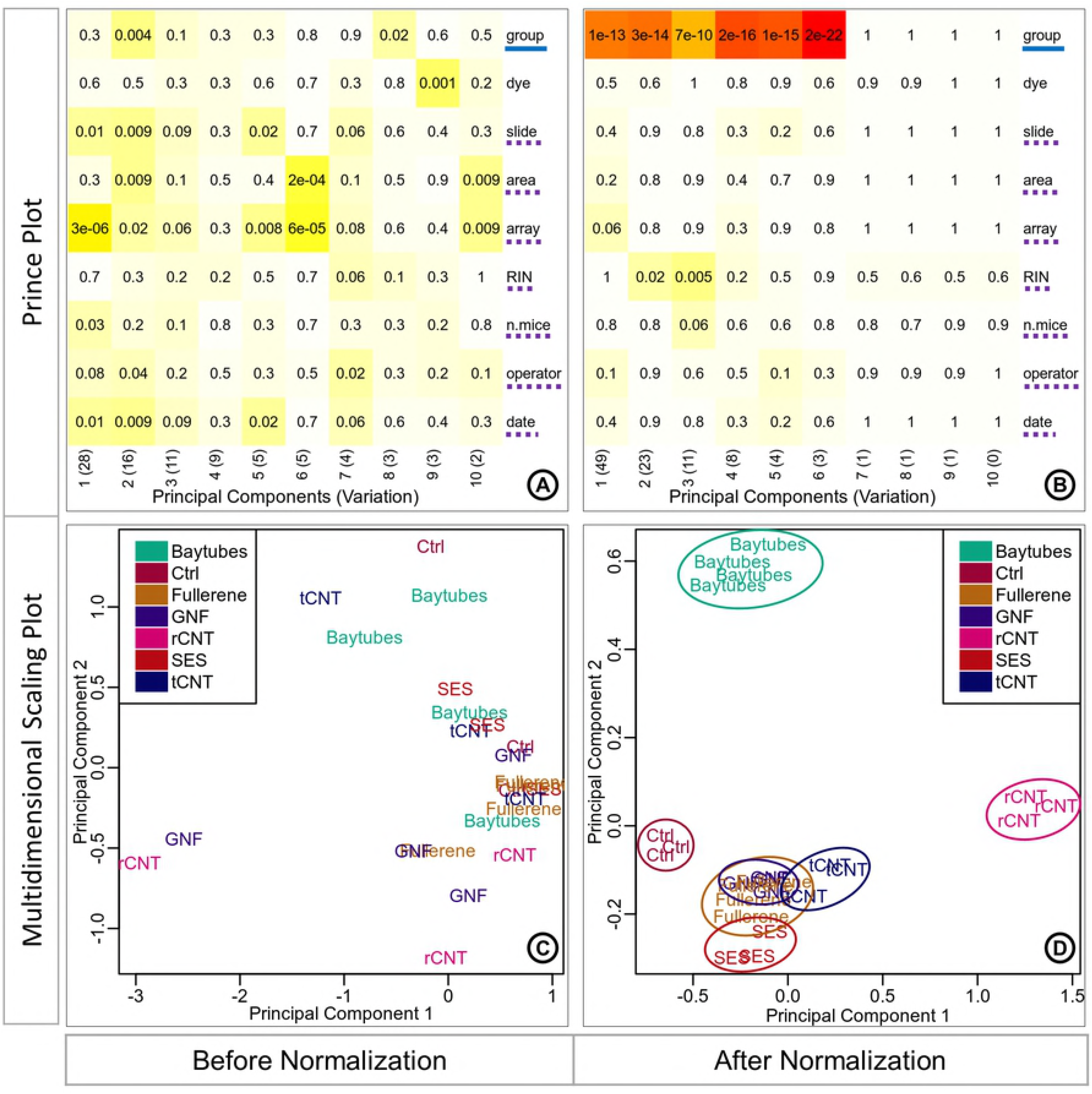
Known batch correction effect. Prince plots (A; B), and PCA (Principal Component Analysis) plots (C, D) before and after batch correction. The prince plots (A, B) report the association between the first ten principal components of the data and the annotation variables, principal components are represented in columns, and the sample annotations are reported in rows. The cell text labels, cell background color, and row label notations with solid blue underline and dotted purple underline are the same as Fig4A. PCA plots (C, D) report the relationship between the samples as a scatter plot. The circular outlines in the PCA plot after correction (D) are provided for better visibility of sample grouping.

The hidden sources of variation can be identified as surrogate variables by using sva method. The attributes of these surrogate variables must be inspected before taking any decisive action. These sources of hidden variation can represent technical batch information or biological information such as sub-types. Thus, care must be taken to choose the right candidate variables for batch correction. The sva analysis resulted in three surrogate variables that were significantly associated with the first three principal components representing high variation as observed in the prince plot (Fig 6A). These identified surrogate variables show significant association with known batch variables ‘slide’, ‘area’, ‘array’, ‘n.mice’, ‘operator’, and ‘date’ (Fig 6B). It is noticeable that none of these surrogate variables have a significant association with the ‘dye’ variable which can be explained by the fact that ‘dye’ variable is not associated with any principal component representing high variation in the prince plot generated from uncorrected data (Fig 6B).

**Fig 6.**
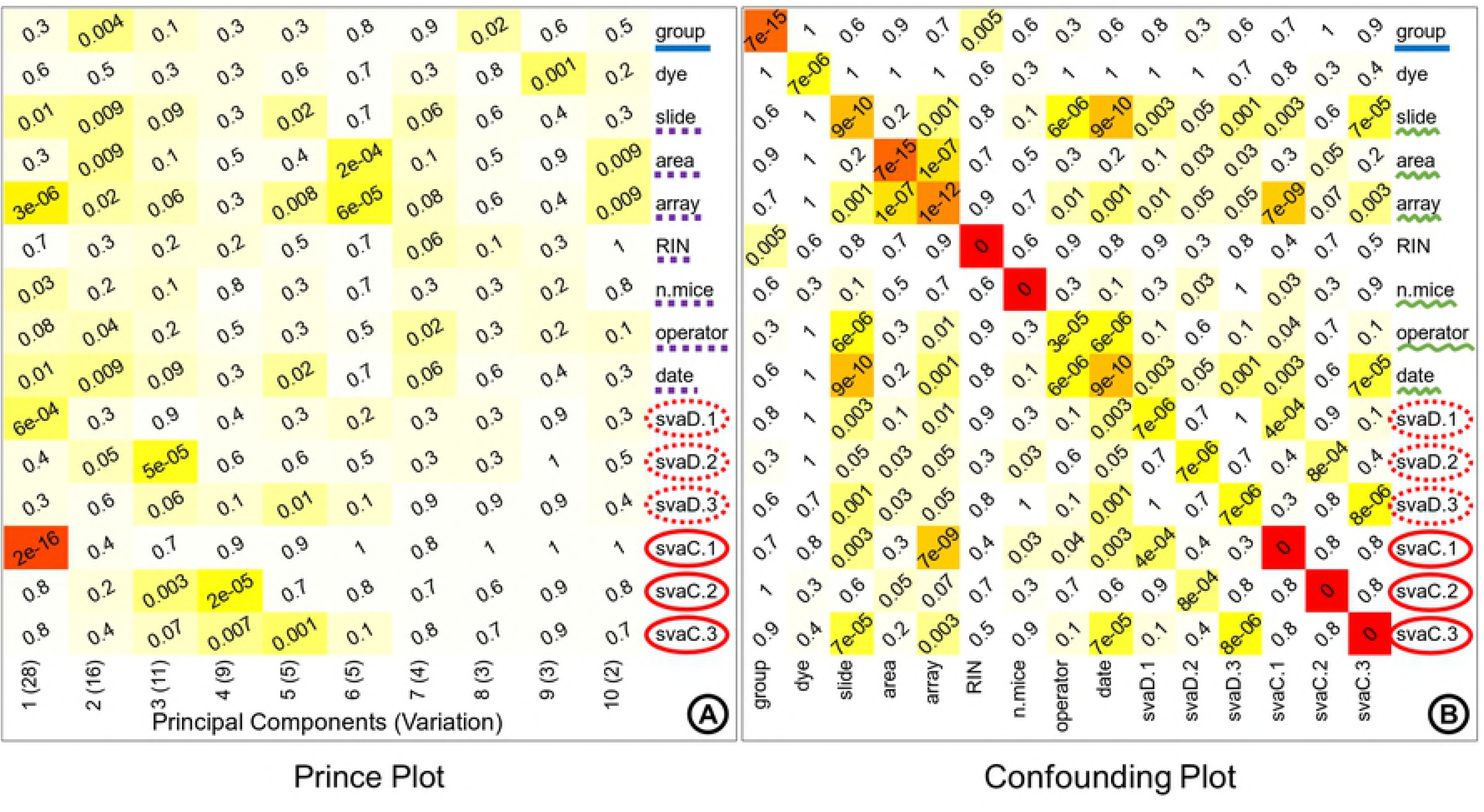
Hidden batch attributes. The surrogate variables identified from sva (surrogate variable analysis) are represented in discretized and the original continuous form. These variables are combined with the known variables to generate a prince plot (A) and confounding plot (B). The prince plot (A) reports the association between the first ten principal components of the data and the annotation variables, principal components are represented in the columns, and the sample annotations are represented in the rows. The cell text labels, cell background color, and row label notations with solid blue underline and dotted purple underline are the same as Fig4A. The confounding plot (B) reports the association between all pairs of annotation variables; annotation variables are represented in both rows and columns. The cell text labels, cell background color, and row label notations with solid blue underline are the same as Fig4A. The row labels underlined with squiggly green line represent known variables associated with surrogate variables. The row labels circled by dotted red outline in (A) and (B) represent the discretized surrogate variables. The row labels circled by solid red outline in (A) and (B) represent the continuous surrogate variables.

The discretized surrogate variables were associated with the first three principal components in the prince plot generated from data before batch correction (Fig 7A). The variable of interest ‘group’ was significantly associated (p-value: 0.004) with the second principal component. While the surrogate variables ‘svaD.1’ was significantly associated with the first principal component (p-value: 6e-04), ‘svaD.2’ was significantly associated with the second and third principal components (p-value: 0.05, and 5e-05, respectively), and ‘svaD.3’ was significantly associated with the third principal component (p-value: 0.06). In the prince plot generated from the data after batch correction (Fig 7B), the variable of interest ‘group’ is now observed to have a high association with the first principal component (p-value: 3e-07). The discretized surrogate variables ‘svaD.1’, ‘svaD.2’, and ‘svaD.3’ are no longer associated with the first three principal components representing high variation. The removal of hidden batch effects with surrogate variables removes artefactual technical variation from the data, thus revealing the true variation signal of the variable of interest. The PCA plot generated from the data before batch correction (Fig 7C) displays samples scattered across the projected components with no obvious grouping of samples by the variable of interest ‘group’. The grouping of samples in the PCA from the corrected data (Fig 7D) is more discrete than the uncorrected data, but the separation between the groups (intergroup distances) is not very clear, only ‘rCNT’ and ‘Ctrl’ show visibly distinct separation (Fig 7D), while the samples from other groups are much closer. The tighter packed grouping of samples can be observed by the circular outlines drawn around the groups in Fig 7D.

**Fig 7.**
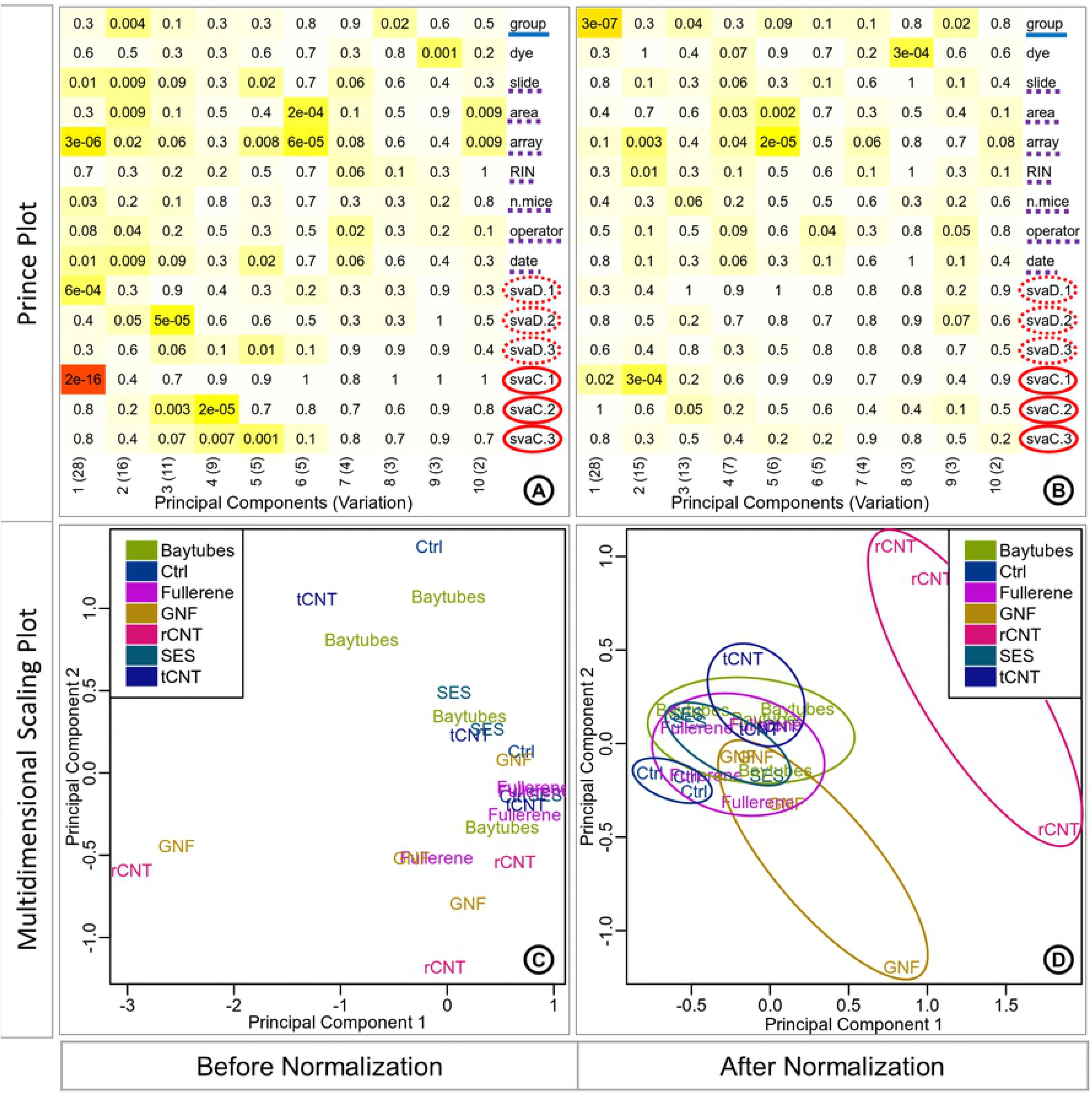
Hidden batch removal effect. Prince plots (A; B), and PCA (principle component analysis) plots (C, D) before and after batch correction. The prince plots (A, B) report the association between the first ten principal components of the data and the annotation variables, principal components are represented in columns, and the sample annotations are represented in rows. The cell text labels and background color notations are the same as Fig4A. The row label notations with solid blue underline, dotted purple underline, dotted red outline circle, and solid red outline circle are the same as Fig6A. The PCA plots (C, D) report the relationship between the samples as a scatter plot. The circular outlines in the PCA plot after correction (D) are provided for better visibility of sample grouping.

The differential expression analysis is performed by specifying the limma linear model parameters in the eUTOPIA interface. Here, the ‘group’ annotation variable is specified as the ‘variable of interest’. The known batches and the surrogate variables are instead specified as covariates for the corrected data. The contrasts are defined to compare each nanomaterial ‘Baytubes’, ‘Fullerene’, ‘GNF’ (Graphite Nano Fibers), ‘rCNT’, ‘SES’, and ‘tCNT’ to the control ‘Ctrl’ samples. The differential expression analysis results are filtered by logFC cutoff threshold 1 and P.Value cutoff threshold 0.05. Differential expression results are explored by the medium of plots (Fig 8) to classify contrasts and conditions by expression patterns. The intersection of differentially expressed gene sets represented as an UpSet plot (Fig 8A) can be used to infer that rCNT samples have overall larger variation contrast against the control (Ctrl) samples, while the rest of the nanomaterials have much smaller and comparable variation contrasts. There is higher overlap between rCNT and tCNT sets with larger intersection sizes and multiple intersections. Nanomaterial specific genes are substantial in rCNT, GNF, Baytubes, and tCNT while specific genes are not reported for SES and Fullerene because of their smaller values. The distribution of differential genes from each contrast is represented as a volcano plot (Fig 8B) which can be used to infer that the genes in this particular contrast/comparison are more inclined towards overexpression with higher significance values while the underexpressed genes are fewer and have lower significance values. The expression pattern of top differentially expressed genes is represented as a Heatmap (Fig 8C), which can be used to infer that there are two major parent clusters of samples. The first parent cluster contains a child cluster of Fullerene (hollow carbon sphere) and GNF (long rigid carbon fiber), which is further clustered with rCNT (long rigid multi-walled CNT). The second parent cluster contains two child clusters, control samples (Ctrl) cluster together with Baytube (short tangled multi-walled CNT) to form the first child cluster, SES (short rigid multi-walled CNT) and tCNT (long tangled multi-walled CNT) cluster together to form the second child cluster. The nanomaterials in the first parent cluster have a larger diameter and smaller surface area compared to the nanomaterials in the second parent cluster. The expression pattern of the selected top differentially expressed genes align very well with the nanomaterial intrinsic properties. The intrinsic properties of the nanomaterials are provided in S3 File.

**Fig 8.**
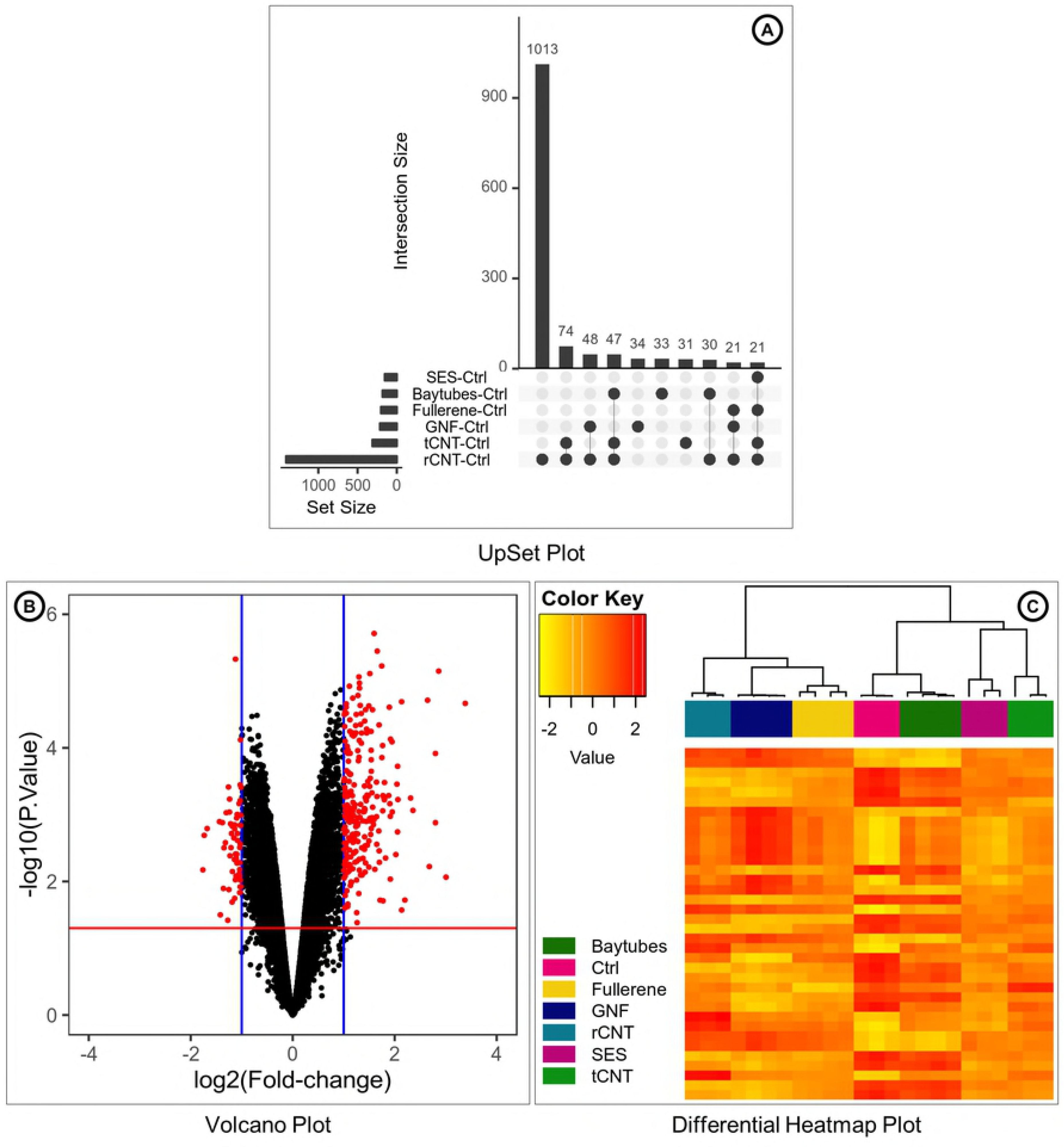
Differential expression summary. Selection of plots is reported by eUTOPIA for the differential expression analysis results. The UpSet plot (A) represents the intersection between the sets of differentially expressed genes from various comparisons/contrasts in the limma setup. The vertical bar plot reports the intersection size, the dot plot reports the set participation in the intersection, and the horizontal bar plot reports the set sizes. The volcano plot (B) represents the distribution of differentially expressed genes by log2(Fold Change) and -log10(P.value) as a scatter plot. The log2(Fold Change) values are plotted on the x-axis, the range of values is from the negative logFC to positive logFC with zero in the middle. The blue vertical margins on the plot represent the user specified logFC cutoff. The -log10(P.Value) values are plotted on the y-axis, the red horizontal margin represents the user specified P.Value cutoff. The plotted values outside the cutoff margins are colored red to highlight the significant differentially expressed features. The heatmap (C) represents the top differentially expressed features. The samples are reported in rows, and the features are reported in columns. The log-transformed expression values are represented by the color as per the ‘Color Key’. The clustering of samples based on the expression distribution of the reported features is presented on the top of the heatmap. The color blocks below the hierarchical cluster represent the sample group annotation.

## Results and discussion

eUTOPIA allows users to reliably process microarray data and generate visual interpretations of the results seamlessly *via* a simple yet intuitive interface. It is focused on the preprocessing of microarray data, thus providing an agile and robust alternative to more comprehensive tools. Both commercially available and free software for microarray data preprocessing either provide limited methodological options or force the user to design an appropriate pipeline from the tools provided. This can be a daunting task for researchers with limited experience in omics data analysis. eUTOPIA is designed to balance both reliability and flexibility.

The case study showcases eUTOPIA’s features that enable the user to perform array data preprocessing and analysis. eUTOPIA’s guided workflow helps to easily preprocess the data, identify sources of unwanted variation, remove them and evaluate the sanity of corrected data. The graphical user interface allowed the ease of defining a linear model to test the data and the wide choice of provided dynamic plots help to classify and characterize conditions by sets of differential features and expression patterns.

We compared eUTOPIA against a set of microarray analysis tools that are free for academic use and have a graphical interface, namely AGA [14], shinyMethyl, MeV [15], O-miner [16], Chipster [17], and Babelomics [18]. The comparison table (S4 File) evaluates the tools over a list of implemented analytical steps and supported data platforms. One major feature that is not supported by most of the compared tools is the *Batch Correction* of *Known Variables* and more noticeably of *Surrogate Variables*. eUTOPIA’s analysis workflow integrates the visual representation of sample annotation and principal components of variation to identify batch variables, along with the ability to perform correction of batch effects seamlessly. Furthermore, it is the only tool that incorporates the surrogate variable (hidden batch) identification, visualization, and correction. This process of batch identification and correction is of extreme importance for microarray analysis because it can help to isolate technical noise from the biological signal. In this comparison, Chipster has the most comprehensive toolbox; it provides the most features that even extends beyond microarray analysis. However, these features are presented as separate tools with no specific workflows and guidelines for choosing the most optimal set of tools. This can pose a challenge to the users in need of designing analysis workflows by combining these tools appropriately. In contrast, eUTOPIA incorporates a specific set of tools in a streamlined workflow to ensure intuitiveness and ease of use. eUTOPIA does not impose on the user the technical challenges of workflow design thus allowing to focus more on the biological aspects of the data and results.

## Availability

eUTOPIA is free for academic use and can be obtained from the GitHub repository https://github.com/Greco-Lab/eUTOPIA.

## Acknowledgments

We would like to thank Nanna Fyhrquist (University of Helsinki, Karolinska Institute) and Marit Ilves (University of Helsinki) for providing the valuable user feedback during the development process.

## Authors Contributions

Conceptualization: VF, DG. Funding acquisition: HA, BG. Methodology: VSM, GS, VF. Software: VSM, AS, VF. Supervision: HA, VF, DG. Validation: GS, AS, HA. Visualization: VSM. Writing - Original Draft Preparation: VSM. Writing - Review & Editing: VSM, GS, PASK, AS, HA, VF, DG.

## Supporting information

**S1 File. eUTOPIA user manual.** eUTOPIA user manual with sample analysis.

**S2 File. Phenotype table.** This phenotype table contains information about the microarray (dye, slide, area, array, file), sample identification and grouping (SampleID, group), quality estimation of extracted RNA (RIN, Qubit_conc, dye_conc, dye_activity), technical information (n.mice), and information associated with microarray experiments (operator, date).

**S3 File. Carbon nanomaterial intrinsic properties.** This table contains the description and intrinsic properties of the carbon nanomaterials used in the case study. General description of the nanomaterial and nanomaterial shape are provided along with the average values of the intrinsic properties’ length, diameter, and surface area. Aspect ratio is the average length divided by the average diameter

**S4 File. Comparison table.** Comparison between eUTOPIA and other open source tools with the same scope.

